# *Dictyostelium discoideum* chemotaxis is altered by hypoxia to orient streaming toward higher oxygen level independently of aerotaxis

**DOI:** 10.1101/2025.08.22.671833

**Authors:** Satomi Hirose, Julie Hesnard, Kenichi Funamoto, Jean-Paul Rieu, Christophe Anjard

**Affiliations:** Graduate School of Biomedical Engineering, Tohoku University, Sendai, Japan; Institute of Fluid Science, Tohoku University, Sendai, Japan; Université Claude Bernard Lyon 1, CNRS, Institut Lumière Matière, UMR5306, Villeurbanne, France

## Abstract

**Background:** *Dictyostelium discoideum* (Dd) exhibits a unique life cycle marked by its transition from single-cell amoeboid movement to multicellular development in response to environmental stimuli. While its chemotactic response to cyclic adenosine monophosphate (cAMP) has been extensively studied, experiments are usually carried out in a high oxygen environment (21% O_2_, also called normoxia) that might not reflect the natural condition for a soil amoeba as hypoxia is common underground. Our recent research has unveiled a novel phenomenon termed aerotaxis, wherein Dd cells migrate towards oxygen-rich environments under low oxygen (hypoxia).

**Results:** We tested Dd response when submitted to two different stress sources: starvation and hypoxia. While both stresses induce a motile response, namely chemotaxis and aerotaxis, we were able to decoupled both type of responses to explore potential shared mechanisms. Importantly, aerotaxis appears to be operated independently of known chemotactic pathways, demonstrating unique signaling pathways and cellular responses. Aerotaxis contributes to Dd developmental processes by letting cells escape from acute hypoxia. Even though chemotactic response can occur at less than 2% O_2_, it does not lead to well organized or stable streaming. Furthermore, in an oxygen gradient, aggregation center for chemotactic response appears preferentially at higher O_2_ level. RT-qPCR analysis show that hypoxia only slightly reduces the expression of genes required for chemotactic response, suggesting that the bias toward high O_2_ might occur at another level. Reoxygenation of cells that have been starved in hypoxic condition for 18h allow rapid aggregation but require *de novo* protein synthesis.

**Conclusion:** Aerotaxis and chemotaxis mechanisms do not interact directly. However, when cells are exposed to both hypoxia and starvation, both mechanisms can be combined to direct the migration of cells toward place with higher oxygen level. As hypoxia is frequent in the soil where Dd usually grow, formation of aggregation centers at, or close to the soil surface, where oxygen is abundant, will be advantageous. Low O_2_ level does not preclude cells to participate to streaming but seems to reduce or delay their ability to produce stable aggregation centers, resulting into bias favoring centers that form at higher oxygen level.

## Background

The social amoeba *Dictyostelium discoideum* (Dd) is an obligate aerobic organism that lives in the soil, feeding on bacteria. Dd exhibits a distinctive life cycle, transitioning between unicellular and multicellular forms. In nutrient-rich conditions, Dd lives as individual amoeboid cells. Upon starvation, however, the cells initiate a developmental program involving aggregation and differentiation, ultimately leading to the formation of fruiting bodies that ensure spore dispersal (1,2). This transition relies on the capacity of Dd to sense and respond to a wide range of environmental cues, including chemicals, temperature (3), light (4,5), electric fields (6,7), and oxygen (O_2_) (8). A central component of this developmental response is chemotaxis toward cyclic adenosine monophosphate (cAMP), a key signaling molecule that mediates long-range intercellular communication and directs coordinated migration (9,10). Chemotactic cells polarize and elongate along cAMP gradients, relaying the signal to their neighbors and organizing into head-to-tail streams that enhance collective motility (11,12). Although the initial positioning of aggregation centers appears largely random, competition and fusion events eventually lead to evenly spaced territories, each comprising approximately 10^5^ cells. At the molecular level, starvation triggers a rapid and temporally regulated transcriptional reprogramming (13), inducing genes involved in signal production and sensing, such as adenylyl cyclases responsible for cAMP synthesis, and cAMP receptors (CARs) required for gradient detection. As development proceeds, Protein Kinase A (PKA) becomes activated and promotes the transition to later stages, including terminal differentiation. The developmental process is also closely associated with the induction of several cell adhesion proteins. Notably, the contact site A glycoprotein (CsaA) and the calcium-dependent adhesion molecule (CadA) are expressed shortly after the onset of development (14,15). In contrast, the transition from the unicellular to the multicellular stage is governed by the polymorphic adhesion molecules TgrB1 and TgrC1, which also play a central role in allorecognition (16). Although the molecular basis of this coordinated motility is well understood, the initial establishment and maintenance of aggregation centers remain less clear (17).

In recent years, using a double-layer microfluidic device able to control precisely -O_2_ gradients, we found that *Dd* single cells in a nutrient medium detect and respond to O_2_ gradients below ∼2% O_2_, exhibiting both directional migration toward higher O_2_ (aerotaxis) and increased speed in low-O_2_ (aerokinesis) (18,19). Using a simple confinement assay consisting in covering a spot of *Dd* cells with a cover glass, we also showed that O_2_ consumption induces self-generated O_2_ gradients with a concentration at the spot center below 0.1% O_2_ and around 0.5% O_2_ at the edges (18). This O_2_ gradient combined with aerotaxis triggers the formation of a robust migrating dense cell ring (18), as well observed in an independent study (20). However, the molecular mechanisms behind the O_2_ sensing in Dd cells and how it affects motility remain unknown. Our previous work excluded the involvement of PhyA, the Dd homolog of mammalian prolyl hydroxylase (PHD) enzymes, which regulate the hypoxia-inducible factor (HIF) modification and trigger the transcriptional response to hypoxia (19). Moreover, we showed that the aerotactic migration and O_2_ sensing of Dd cells are not hampered by the ROS/RNS and inhibition of mitochondrial functions (19).

As starvation and lack of O_2_ (hypoxia) are common stresses for a soil dwelling micro-organism, we explored how *Dd* respond when subjected to both. To this end, we first conducted experiments under controlled O_2_ gradients using a microfluidic system, and employed chemical treatments to decouple aerotactic responses from starvation-induced signalling and cAMP-mediated chemotaxis. We further explored how O_2_ availability modulates chemotaxis by analyzing the spatial patterns of aggregation under different O_2_ concentrations. To complement these observations, we performed RT-qPCRs to assess O_2_-dependent gene expression changes. Finally, we evaluated the effect of reoxygenation on streaming formation and investigated the role of *de novo* protein synthesis in this process.

## Methods

### Cell culture

The wild types AX2 and AX3 of Dd cells were provided by the National BioResource Project (NBRP Nenkin, Japan). The cAR1 and cAR3 null mutant RI9 cell line was a kind gift from Prof. Satoshi Sawai from Tokyo University, Japan. AX2 was used for all experiments unless otherwise mentioned. Cells were cultured axenically with HL5 medium (HLF2; Formedium, United Kingdom) containing 10 g/L glucose and 10 U/ml penicillin-streptomycin (P/S) (15140-122; Gibco, United States) on cell culture dishes at 22°C or in shacking culture at 180 rpm. For starvation conditions, cells were cultured in phosphate buffer (PB) solution (12 mM KH_2_PO_4_, 8 mM Na_2_HPO_4_, pH 6.5) or the glucose-free HL5 medium.

### Oxygen gradients experiment in microfluidic devices

Experiments under O_2_ control were done in the microfluidic device as we previously reported (19,21) and described in Supplementary Figure S1. Briefly, the device has two gas channels to supply gas mixtures at predefined O_2_ concentrations, above the media channels where Dd cells were seeded. An O_2_ gradient was generated in the media channel by supplying the gas mixtures at 0% and 21% O_2_ balancing with nitrogen at 30 mL/min (Figure S1B). The gradient was calculated using COMSOL Multiphysics (ver. 5.5, COMSOL AB, Sweden). For the detailed features and performance of the device, please see our previous publication (21).

Dd cells were harvested from the culture dishes and introduced into the media channels in the microfluidic device with the HL5 culture medium. Cell density was 2 × 10^6^ cells/mL and 1 × 10^7^ cells/mL for a short-term experiment to observe aerotaxis and a long-term experiment to observe cell aggregation under starvation, respectively. After incubation for 20 min to adhere Dd cells to the bottom surface of the media channels, the HL5 culture medium was replaced by fresh HL5 media or PB solution for starvation conditions. Caffeine (C2042; Tokyo Chemical Industry, Japan) was added at a final concentration of 3 mM while thapsigargin (029-17281; Fujifilm Wako Chemicals, Japan) was to added to the cell culture media at 2 or 5 µM as indicated. For accurate comparisons, one of the three media channels in each device were used for experiment under the control condition, while the other channels were used simulaneously for experiments under different conditions. The device was placed in a stage incubator (TP-CHSQ-C, Tokai Hit, Japan) mounted on an inverted microscope (IX83, Olympus, Japan), and controlled at 22°C. Just after gas mixtures were supplied to the gas channels, time-lapse imaging was started at a 5-min frame rate. As previously described, we calculated the strength of the aerotaxis with the aerotactic index (*AI*), defined as the ratio between the distance migrated in the direction along the O_2_ gradient and the total length of cell trajectory (Figures S1C-E), thus in the same way than chemotactic index was calculated (19). Moreover, the position of initial aggregation centers distribution was investigated manually as it corresponded to the initial point of convergence of chemotactic cells.

### Streaming assay in homogenous hypoxic and normoxic conditions

The assay is adapted from standard streaming assay to allow multiposition videomicroscopy simultaneously in a homemade hypoxic chamber as well as in ambiant conditions. AX2 cells were washed once with developmental buffer (DB) solution (20 mM phosphate buffer, pH 6.5; 1 mM CaCl_2_; 2 mM MgSO_4_) and then seeded in two different 6 well plates (at 1 or 2 × 10^6^ cells per well). After 30 min, 2 ml of light paraffin oil (VWR, CAS:8012-9S-1VWR) was slowly added on the top of each well to allow imaging for over 24 h without evaporation or condensation. One plate was incubated in the hypoxic chamber preconditioned with the different O_2_ level (1%, 2% or 4% O_2_) while the other was maintained under normoxic (21% O_2_) atmospheric condition. The O_2_ concentration in the hypoxic chamber was monitored with an optical fiber probe coupled with its respirometer (OXROB3 robust probe and Firesting oximeter, Pyroscience, Germany). Both plates were positioned on a motorized videomicroscopy platform allowing imaging of both hypoxic and control conditions. This platform, controlled by LAS X software (Leica, Germany), includes a binocular microscope (MZ16, Leica Germany), a motorized stage (XY-Scanning stage 150 x 100 mm, Marzhauser, Germany), a CCD camera (DMC2900, Leica, Germany) and a transmitted light base (TL300, Leica, Germany) to obtain dark field images. Images were taken at a 10 min frame rate and were subsequently analyzed and filtered using basic ImageJ functions. Image substraction using a one-frame difference was performed to enhance dark field contrast.

Reoxygenation was induced by opening the hypoxic chamber and stopping the gas flow. Cycloheximide (C7698, Sigma-Aldrich) from a 200 mM stock in DMSO was added at 400 µM 30 minutes prior reoxygenation to inhibit protein synthesis. DMSO (D8418, Sigma-Aldrich) alone was used in the control condition. Cycloheximide was added approximately 18 h after the cells were placed under 1% O_2_, as preliminary experiments showed that reoxygenation after 24 h of hypoxia sometimes resulted in poor or absent aggregation, even in the absence of treatment. The experiment was performed three times.

### RT-q-PCR

A total of 2 × 10^6^cells per well were washed once with DB and then seeded in 35 mm single well plates. Two plates were incubated in our hypoxic chamber preconditioned with 1% O_2_, while the other two were maintained under normoxic (21% O_2_) atmospheric conditions. The O_2_ concentration was monitored with the above mentioned optical fiber probe coupled with its respirometer. Cells were harvested after 6 and 24 hours of incubation in both O_2_ conditions. Additionally, 2 × 10^6^cells were collected directly from HL5 culture medium to serve as the standard condition t0.

Total RNA was extracted (RNeasy Plus Mini Kit, 74134, Qiagen, Germany), quantified and adjusted at 500 ng to synthesize cDNA (First Strand cDNA Synthesis Kit for RTPCR,11483188001, Roche, Switzerland) using random primers at 250ng/μl (DyNAmo Flash SYBR Green qPCR Kit, F415L, Thermo Fisher Scientific, USA). cDNAs were diluted at 1/10 to perform qPCR (Thermocycler BIOER qPCR LINEGENE 9600 PLUS), using the following primers: acaA-F 5’-CCAATGTTTAATGATCATGC-3’, acaA-R 5’-ATGATACACCTCCTAAATGTG-3’, pkaC-F 5’-GTACTGGAACATTTGGAAAG-3’, pkaC-R 5’-GAATGGATGAGAGAATTGAT-3’, tgrB1-F 5’- GGAGTGGTTACTATCAATGGTC-3’, tgrB1-R 5’-GGTTTAGTGCCAGATGTGTAG-3’, cadA-F 5’-GGCTCAAGGCAGTACAAACA-3’, cadA-R 5’-GGTCATTTCATATGAACCAGC-3’, csaA-F 5’CGCATCGTTACAATGCCAATT-3’, csaA-R 5’-AAGACCATTACCACCTGTAGA-3’, rnlA-F 5’-GCACCTCGATGTCGGCTTAA-3’, rnlA-R 5’-CACCCCAACCCTTGGAAACT-3’ cxgs-F 5’-AAGTTGTTAAATCTCAACTC-3’, cxgS-R 5’-TTTATCAACGCCATATTTAA-3’.

The experiments were performed in triplicate for each gene. The relative gene expression was determined using the 2^-ΔΔCt^ method (22) using *rnlA* as a housekeeping gene and the t0 condition as the standard.

## Results

### Aerotaxis is altered under severe starvation

We began our study by evaluating how starvation influences aerotactic behavior using the 2-layer microfluidic device with O_2_ gradient control (supplementary Figure S1). For this purpose, we observed Dd migration directed by positive O_2_ concentration gradient not only in nutrient medium (HL5 cell culture medium), but as well in PB solution without nutrients or glucose-free HL5 cell culture medium, and computed the aerotactic index *AI* (Figure 1A). Under the starvation condition with PB solution, the maximum *AI* was decreased compared to the conditions with HL5 medium and the glucose-free HL5 medium (Figure 1C). Those data suggest that lack of glucose has no effect on aerotaxis while severe starvation decreased cell capacity to migrate towards O_2_.

**Figure 1:**
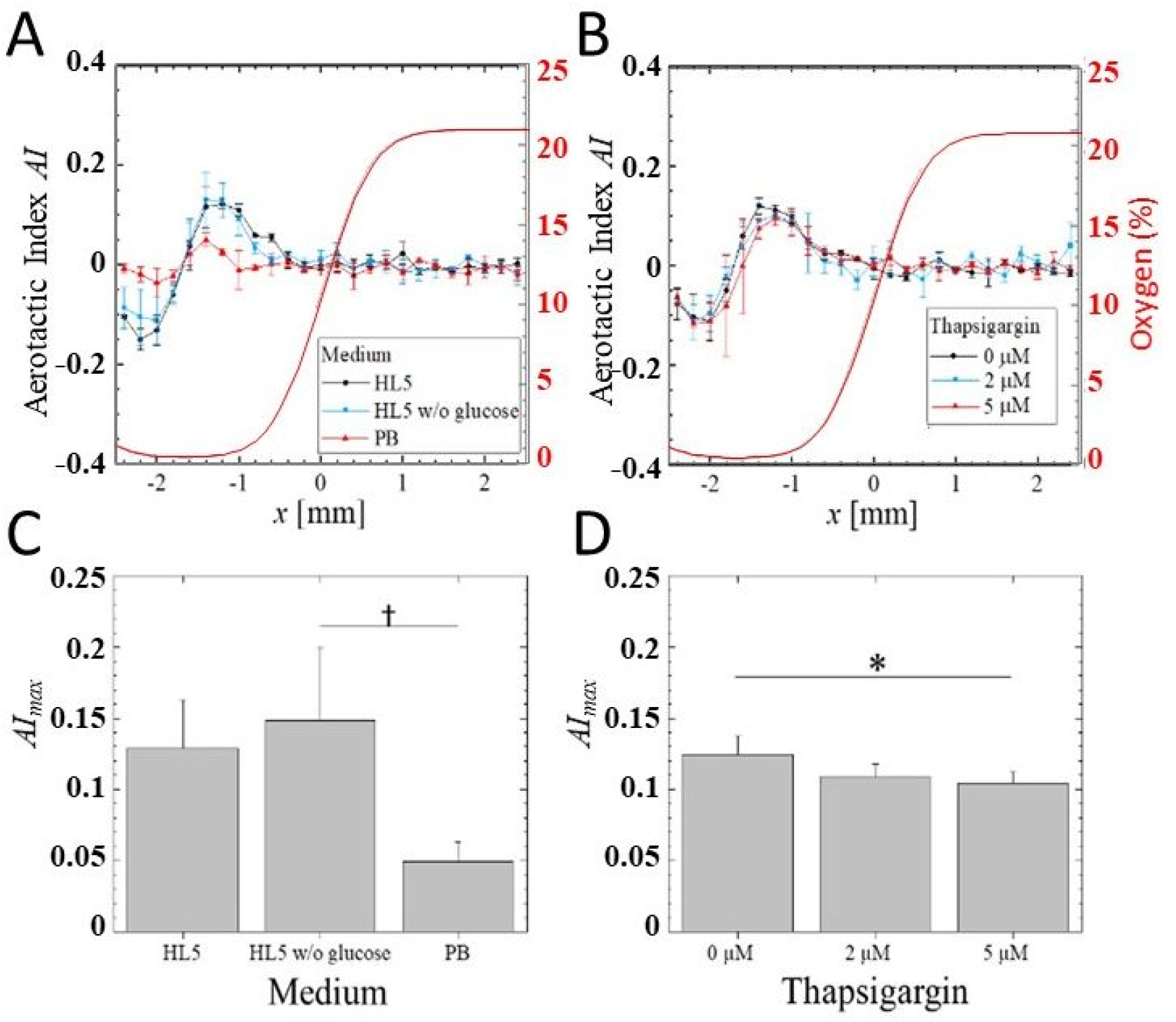
Dd aerotactic index (*AI*) under starvation: A),C) in dif ferent nutritional conditions (HL5, HL5 without glucose and PB solution), B),D) in HL5 with/without thapsigargin. A)-B) Profiles of *AI*, for the various conditions (symbols and lines) along the computed 0-21% O_2_ gradient profile (solid red line) between 1 h and 2 h after the O_2_ concentration gradient was generated. The minimum concentration at x=-1.5mm is ∼0.45% O_2_. C)-D) Maximul *AI* value for each condition. The error bars show the standard deviations of three independent experiments performed in three indipendent devices. For C), Significant differences were assessed by one-way ANOVA followed by Tukey’s post hoc test for multiple comparisons. ^†^*p* < 0.1. For D), Significant differences were assessed by Welch’s t-tests for comparison with the control. \**p* < 0.05.

### Aerotaxis has different mechanisms from known cAMP chemotaxis

Subsequently, we investigated whether aerotaxis shares common signaling mechanisms with established starvation-induced responses and chemotaxis. It has been reported that the starvation response of Dd cells is transduced by an increase in intracellular calcium ions (23) that can also be triggered by addition of thapsigargin, a Ca^2+^ uptake inhibitor of the ER, resulting into enhanced aggregation. Under the microfluidic controlled O_2_ gradient with supplementation of thapsigargin to the HL5 cell culture medium, aerotaxis tended to decrease (Figure 1B, D), showing a similar tendency observed with PB solution. Therefore, signaling pathways relating to starvation might be interfering with the aerotactic response without abolishing it.

Next, association of aerotaxis with cAMP relay was investigated by supplementing 3 mM caffeine, an inhibitor of cAMP relay (24), to the cell culture medium. It is well known that caffeine blocks aggregation of *Dd* under starvation. However, it did not block aerotactic migration under O_2_ gradient (Figure 2A, C). This implies that the cAMP relay is unnecessary for the aerotaxis. To confirm the independence of aerotaxis on cAMP relay, the experiments were also performed with RI9 strain which is deficient in the cAMP receptors cAR1 and cAR3. Lacking those genes, RI9 cells in PB solution are not able to start cAMP relay. We checked that those cells did not show aggregation under starvation (not shown). However, RI9 cells exhibited aerotaxis, indicating the independence of aerotaxis on cAMP relay (Figure 2B, D). Knocking-out cAR can affect quite a lot of cell functions such as actin polymerization and various kinase activities, via signal transduction pathways downstream of cAR (25). This might be the reason for the lower *AI* of RI9 cells as compared to their parent strain KAX3 (Figure 2D). Furthermore, while cell polarity was markedly emphasized during chemotaxis (Figure 2E), for instance by elongated cell shapes, no major change in cell shape and polarity was observed during aerotaxis (Figure 2F). Together, those observations indicate that starvation signaling attenuates aerotactic migration, while cAMP relay is not directly associated with aerotaxis.

**Figure 2:**
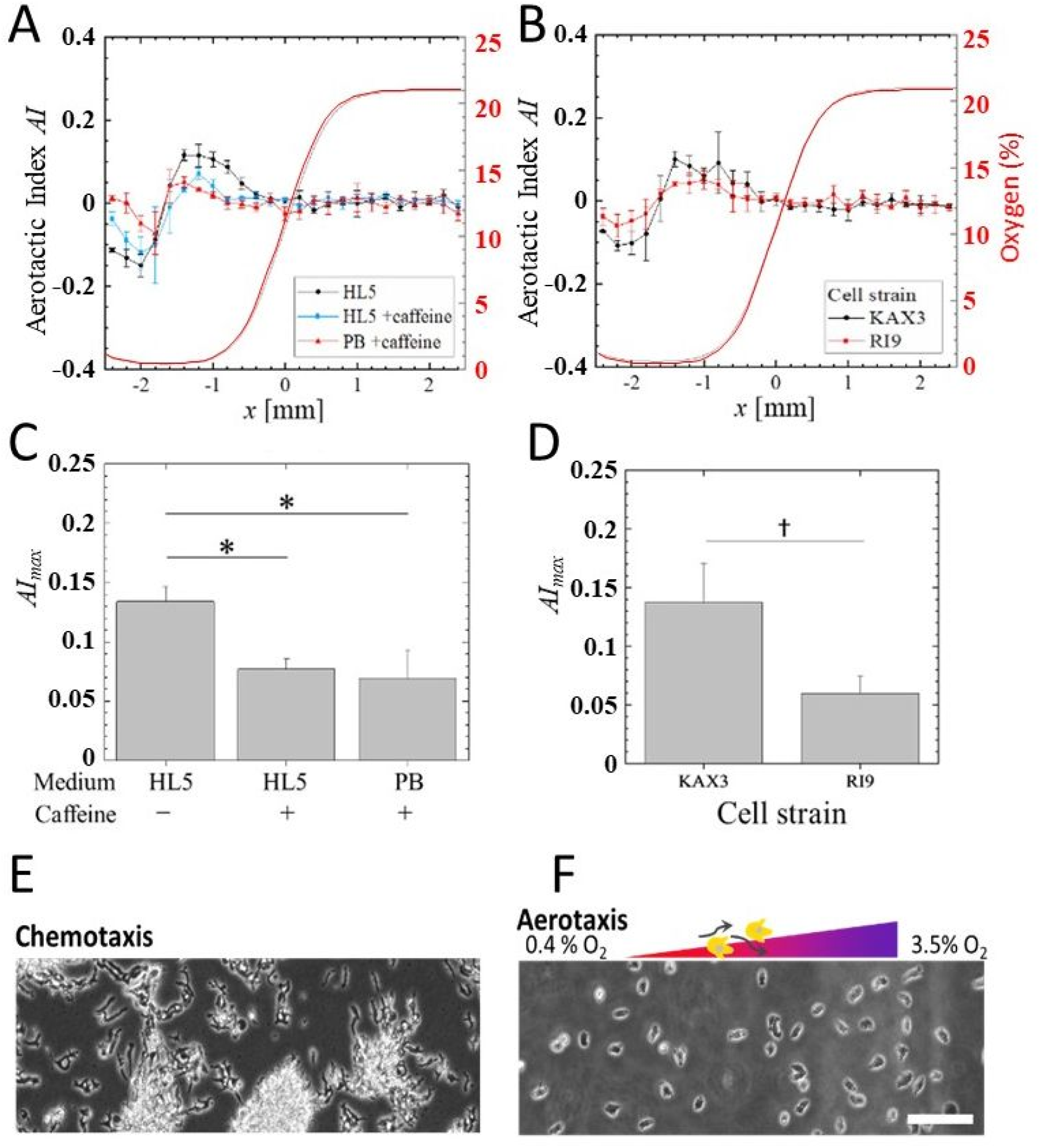
Dd aerotaxis and morphology under chemotactic inhibitor. A)-B) Profiles of *AI*, for the various conditions (symbols and lines) along the computed 0-21% O_2_ gradient profile (solid red line) between 1 h and 2 h after the O_2_ concentration gradient was generated. The minimum concentration at x=-1.5mm is ∼0.45% O_2_. C)-D) Maximul *AI* value for each condition. A)-C) HL5 or PB solution with/without 3 mM caffeine, and B)-D) cAR1^-^/cAR3^-^ deficient RI9 cell line vs its parent strain KAX3. The error bars show the standard deviations of three independent experiments performed in three indipendent devices. For C), Significant differences were assessed by one-way ANOVA followed by Tukey’s post hoc test for multiple comparisons. \**p* < 0.05. For D), Significant differences were assessed by Welch’s t-tests. ^†^*p* < 0.1. E) Cells migrating toward a population by chemotaxis exhibit an elongated shape with high polarity, whereas F) those migrating toward O_2_ rich region by aerotaxis do not exhibit obvious morphological changes (scale bar :100 µm).

### Severe hypoxia disrupts cAMP chemotaxis but not wave propagation

It has been shown above that aerotaxis does not share signaling pathway with chemotaxis or the starvation response, but how is aerotaxis involved in cell development and survival? To investigate the O_2_-dependency of cell aggregation and chemotactic migration, Dd migration under controlled O_2_ conditions was observed under starvation in PB solution for a longer period and at higher cell density (Figure 3 and supplementary movie M1). Under the O_2_ gradient, *Dd* cells showed motility only at O_2_ concentration higher than 0.5% O_2_. As clearly visible in supplementary movie M1, *Dd* cells around the most hypoxic region below that level remained quiescent, they rounded-up and did not migrate, but those located at slightly higher concentration showed aerotaxis and then started to migrate toward aggregation centers like during chemotactic migration in usual normoxic conditions. The initial aggregation centers, defined as the points toward which cells converge, are not distributed randomly but present a clear shift toward high O_2_ concentration, resulting into a chemotactic motility going mostly upward the O_2_ gradient (Figure 3B). In the 7 different usable microfluidic channels, we observed only a single aggregation center occurring under 10% O_2_, most form at higher than 15% O_2_ (Figure 3C). A simple explanation would be that the chemotaxis of cells in hypoxic condition is impaired but careful examination of the movies shows that cells start coordinated motility toward aggregation center at the same time across the O_2_ gradient. Once streaming is more under way, it will split into multiple small aggregates, as observed in normoxic condition.

**Figure 3:**
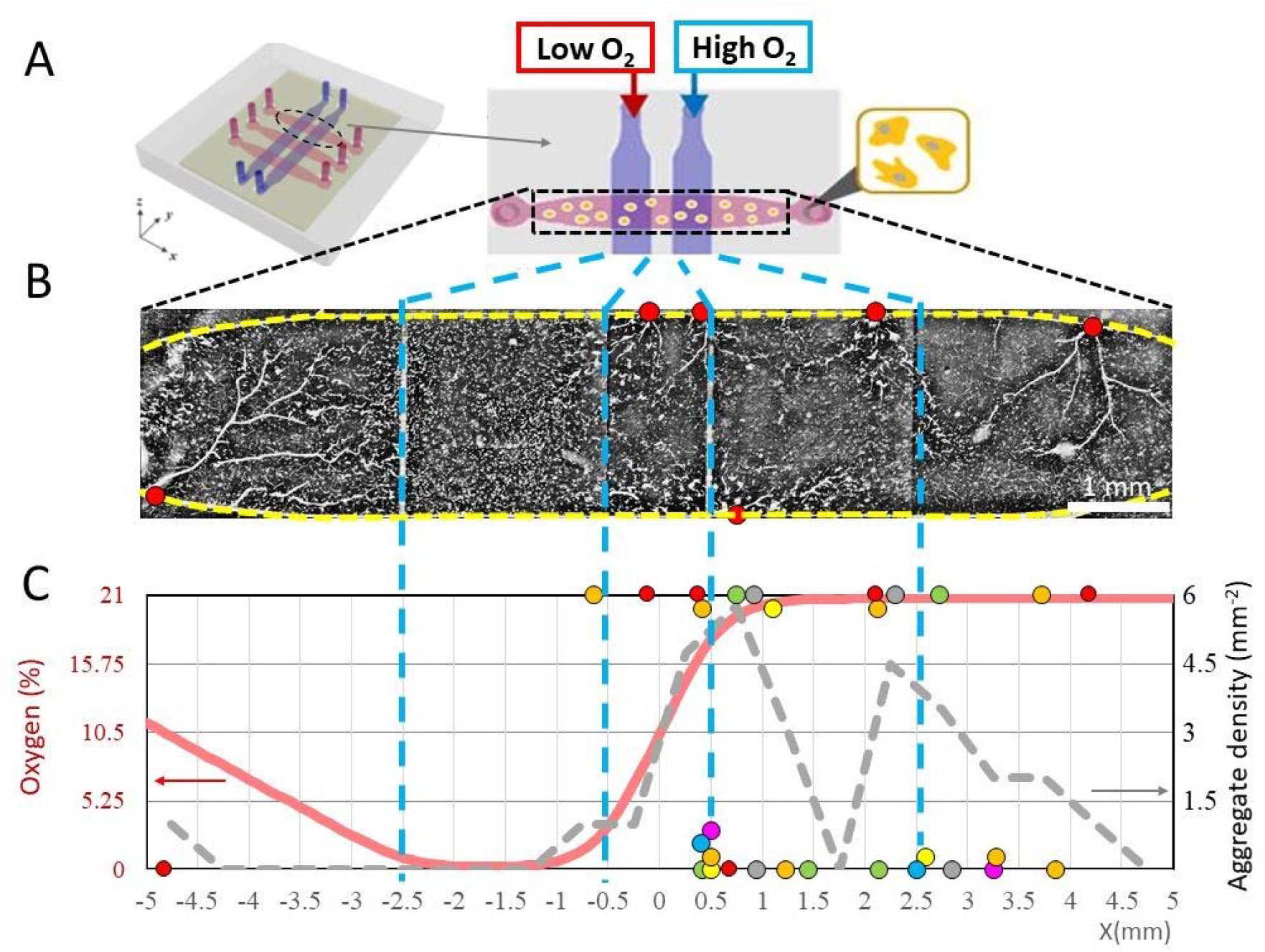
Starvation response of Dd under O_2_ gradient. A) Representation of the microfluidic device and of the observed area in each channel. A multiposition microscope or binocular was used and the full channel images were reconstituted using image J. B) Cell aggregation under starvation in PB buffer under 0-21% O_2_ gradient. The yellow dotted lines highlight the media channel borders. The Dd cells started to chemotax in the region of >0.85 ± 0.01% O_2_). In the representative image, the cells started to stream after 5 h of the change of HL5 cell culture medium to PB buffer and to aggregate toward the locations marked by red circles. C) Plot of the distribution of the aggregation center positions along the microfluidic channel. All 30 observed aggregation centers (n = 7 different channels represented by different colors; N=3 different experiments) are formed on the channel top or bottom border. The distribution is superimposed with the calculated aggregate center density (gary dotted line) along same axis adding both top and bottom points and with the O_2_ profile calculated using COMSOL Multiphysics for a 0-21% O_2_ gradient taking into account the O_2_ consumption by 75,000 cell/cm^2^. The minimum concentration at x=-1.5mm is ∼0.2% O_2_.

To better understand the effect of hypoxia on chemotactic aggregation, time-lapse experiments under submerged uniform hypoxic conditions were performed in 35 mm dishes with a binocular microscope equipped with a transmitted light base in dark field which enables the recording of a large field of view at low zoom, and a motorized stage which enables to perform parallel assays in both a homemade hypoxia chamber and in normoxic control conditions to limit the timing variability. After a few hours of starvation in DB, cell motility appears to increase motility and dark lines corresponding to cell elongating synchronously in response to a cAMP wave are observed using dark field. Those spirals lines start from potential aggregation centers in normoxia that will rapidly lead into organized streaming whatever the O_2_ concentration in the chamber (movies M2-M4). The observed dark lines are somehow less contrasted than the ones previously described (26) with typical dark field setups due to the long distance between the light source and the cells imposed by the size of the hypoxia chamber and the multi-position platform but are nevertheless sufficient to identify the formation of waves especially when using a basic image subtraction method.

Dark bands are forming at 1% O_2_ with a similar timing as in normoxia (Supplementary Figure S2) but don’t result into formation of stable aggregation centers and bands often start to appear in multiple areas in the field of view, resulting into a disorganized pattern (Figure 4A, middle panel) that will sometime fade away (see Movie M2). However, upon reoxygenation, cells will quickly stream and aggregate within 2-3 h (Figure 4A, right panel and movie M2).

**Figure 4:**
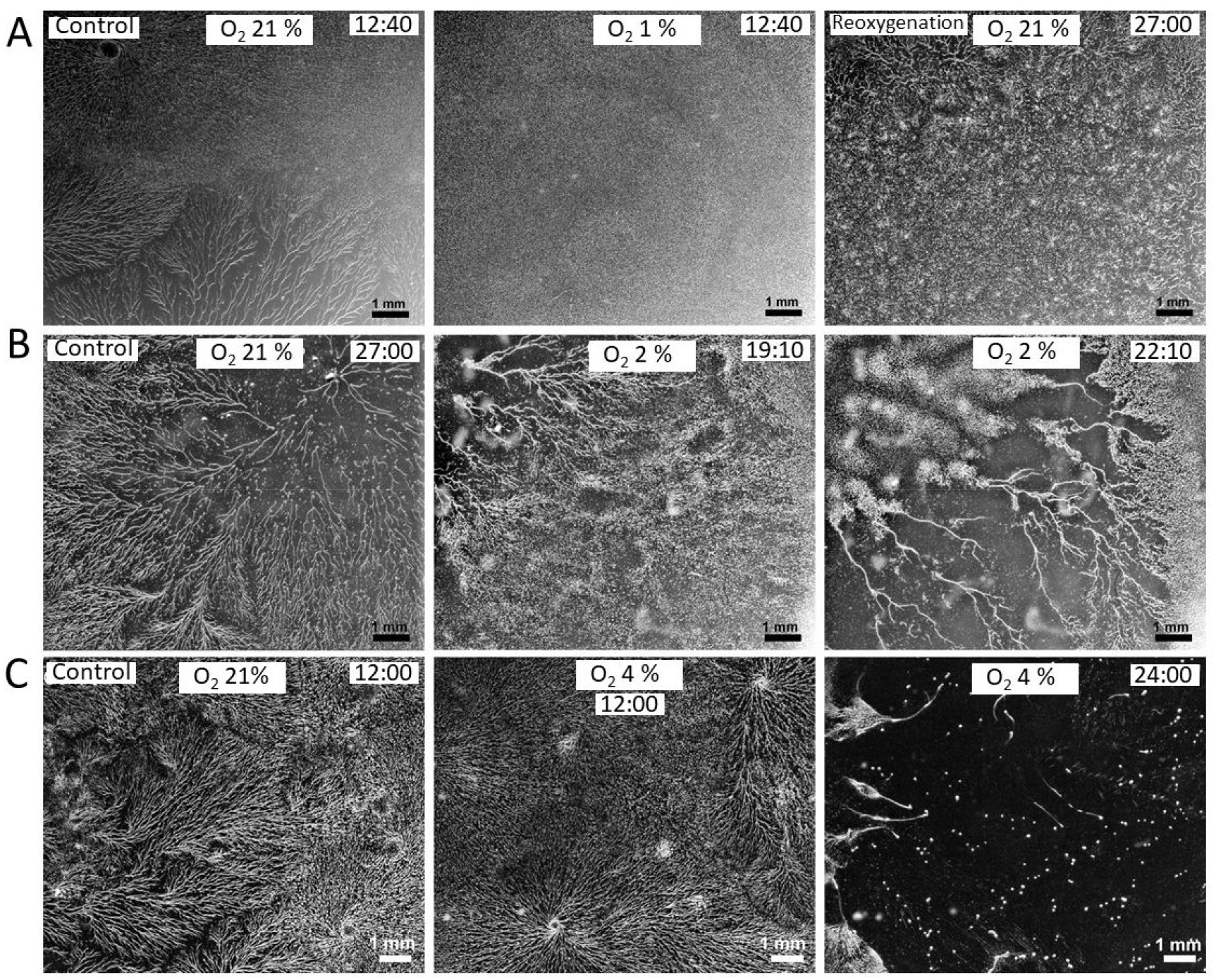
Starvation response under homogenous O_2_ conditions. Representative pictures and movies of aggregation progression for cells in hypoxic (middle and left column) and parallel control normoxic condition of the day (right column). See the high resolution movies M2, M3, M4 for a dynamical vizulation of cAMP induced waves. The experiments under each O_2_ condition were performed at least three times. The scale is indicated on the bottom right while the time from starvation (hour: minutes) is indicated on the top right corner. A) 1% O_2_ vs 21% O_2_ (control). Within 12h40, the control condition shows aggregation streams while the 1% O_2_ condition (middle) present only disorganized waves during 24 h. After 24 h, the sample was reoxygenated and within 3 h, the previously hypoxic sample started to stream and aggregate (right). B) 2% O_2_ vs 21% O_2_ (control). At 2% O_2_, the sample started to stream with some delay with the control but streams quickly dissociatet after their formation and aggregates did not form. C) 4% O_2_ vs 21% O_2_ (control). Streaming did occur at similar time as control and were mostly stable but continued for longer time before forming stable aggregates.

When the same experiment is performed with 2% O_2_ (Supplementary movie M3), the dark bands are followed by organized streaming toward aggregation centers with some slight delay with the normoxia control (Figures 4B, left and middle panels, and Supplementary Figure S2). However, the streams quickly dissociate as individual cells move away from each other at high speed (Figure 4B, right panel). The dissociation tends to start in one area and to propagate at a speed of about 30 to 40 µm per minute. In the presented movies, dissociation started at 19h after the starvation on the upper left corner and propagated toward the center, reaching its maximum after 23.5h, when stream started to reform on the top left while dissociation continue further down. The new streams, will then dissociate again and again. The amount and timing of dissociation is variable from an assay to the next but seem to spread from a “disaggregation center”.

At 4% O_2_, waves then streams were formed and converged into aggregates almost like in normoxia (Figure 4C, left and middle panels), but in some movies, there was some where dissociation events briefly occurred (Figure 4C right side of right panel) then the cells then quickly rejointed nearby streams and formed stable aggregates, albeit with some delay (see Movie M4).

Taken together, these results suggest that hypoxia does not disrupt cell polarity in response to cAMP waves, but inhibits cell aggregation at 1% O_2_ and interferes with cell–cell association at 2% O_2_. The O_2_ threshold required to sustain stable streaming structures is estimated to be around 4% O_2_ but the gradient assays suggest a bias toward even higher levels.

### Hypoxia does not inhibit aggregation gene expression

RT-PCRs were performed to determine if inefficient or unstable streaming in hypoxic condition is due to poor expression of aggregation specific genes (Table 1). The hypoxia-induced gene *cxgS* (19) was included as a control, as it encodes a facultative subunit of cytochrome c oxidase thought to increase the enzyme’s affinity for O_2_ under low-O_2_ conditions (27). *CxgS* showed 50-100 folds induction at both 1% and 2% O_2_ after 6h and 24h starvation, showing that hypoxia can induce gene expression, even during starvation. On the other hand, this gene was repressed after 24 h of starvation in normoxic condition, thus behaving like a growth specific gene. *acaA*, encoding the adenylyl cyclase A (ACA) was induced 300 folds after 6 h of starvation and even more at 24 h. ACA is a key protein responsible for cAMP production during early development that will allow both further aggregation specific gene induction and the organization of chemotactic response for streaming. While hypoxia somewhat reduced ACA induction to 100-200 folds, it remains within error bar of normoxic condition. The expression of *pkaC*, a crucial protein kinase for induction of most early aggregation specific genes, was also similar in hypoxic and normoxic conditions after 6h of starvation but then decline somewhat at 24h in hypoxia. Since 2% O_2_ resulted into unstable streams of cells, we checked the expression level of adhesion genes that are involved in cell-cell adhesion during aggregation. The RNA level of *cadA* barely changed in any condition while *csaA* was somewhat less induced in hypoxic condition after 6 h of starvation but reach similar level than normoxia at 24 h. Like for other most induced genes we analyzed, the high experimental variability between repeats results into values that were within error bar of each other, meaning that the potential differences were not significant. This was true also for *tgrB*, that was induced 2-9 folds after 6 h of starvation in all conditions and 100-300 folds after 24 h. Together, those analyses show that hypoxia only slightly reduces the expression of genes required for chemotactic response, suggesting that the bias toward high O_2_ might occur at another level.

**Table 1:** Expression level of key genes during starvation in hypoxic condition. Cells were placed in PB under hypoxic (1% or 2% O_2_) or normoxic conditions (21% O_2_). Cells were collected after 6 and 24h for RT-qPCR analysis (N = 3). For each gene, the relative gene expression was determined using *rnlA* as a housekeeping gene and the t0 condition as the standard (culture cells in HL5 under agitation).

| Time [h] | % O<sub>2</sub> | *acaA* | *cadA* | *csaA* | *cxgS* | *pkaC* | *tgrB* |
| --- | --- | --- | --- | --- | --- | --- | --- |
| **6** | **1** | 126 ± 161 | 0,6 ± 0,2 | 24,5 ± 33,9 | 105 ± 64 | 2,1 ± 0,8 | 4,4 ± 3,0 |
|  | **2** | 195 ± 56 | 1,6 ± 1,7 | 46,2 ± 42,6 | 85 ± 28 | 2,2 ± 0,3 | 2,4 ± 0,9 |
|  | **21** | 344 ± 160 | 1,7 ± 0,8 | 191 ± 127 | 0,4 ± 0,4 | 3,1 ± 1,0 | 9,1 ± 4,7 |
| **24** | **1** | 160 ± 104 | 1,0 ± 0,5 | 184 ± 182 | 55 ± 41 | 1,4 ± 0,4 | 96 ± 140 |
|  | **2** | 295 ± 32 | 2,3 ± 1,9 | 350 ± 258 | 74 ± 66 | 0,8 ± 0,6 | 235 ± 161 |
|  | **21** | 534 ± 385 | 0,7 ± 0,6 | 141 ± 89 | 0,1 ± 0,07 | 2,8 ± 0,6 | 291 ± 183 |

### Aggregation upon reoxygenation requires *de novo* protein synthesis

As transcriptional analysis doesn’t reveal any major defects in classical aggregation specific genes, we performed an additional experiment to determine if defects in protein synthesis is responsible for the lack of proper aggregation upon hypoxia (Supplementary movie M5). Cells were staved under normoxia and 1% O_2_ as previously (Figure 5A), but the protein synthesis inhibitor cycloheximide or DMSO was added after 18 h incubation (Figure 5B). By then, aggregation was finished in the normoxic control (Figure 5B, left panel). Reoxygenation was then started 30 min later, resulting into rapid aggregation for the DMSO treated cells but not for the cycloheximide treated one (Figure 5C, central and right panels respectively and movie M5). Those experiments indicate that *de novo* protein synthesis is required for streaming to proceed, but it does not exclude some regulation at transcriptional level.

**Figure 5:**
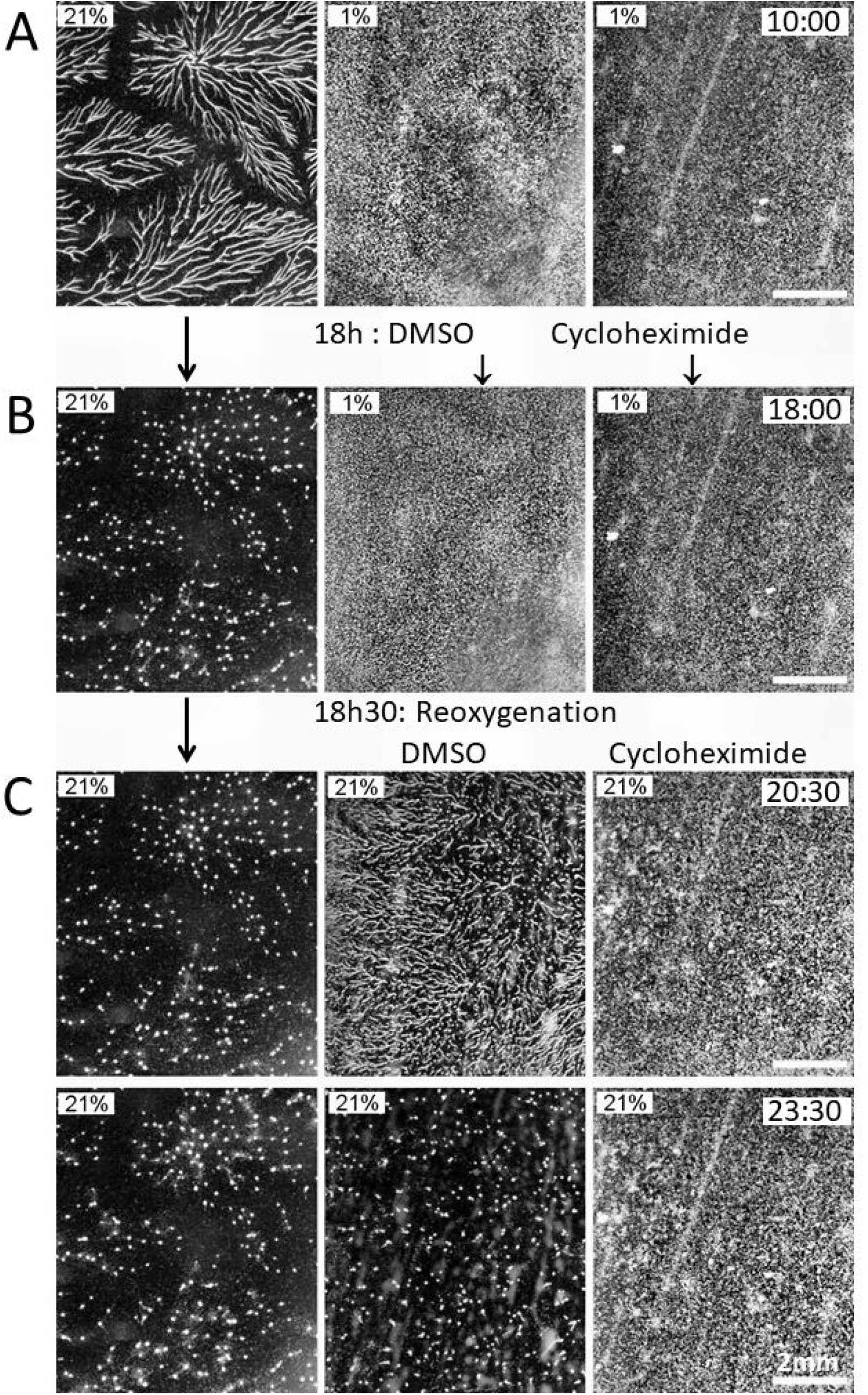
Reoxygenation of starved cells in presence of cycloheximide. A) Cells were incubated at 21% O_2_ (left panel) and 1% O_2_ (middle and right panels) in PB for 18h before addition of DMSO (control) or cycloheximide B) and were reoxygenated at 18h30 by removing the lid of the hypoxic chamber C). The control at 21% O_2_ is also presented. Time of the picture are indicated on the top right, the scales bar correspond to 2 mm.

## Discussion

*Dictyostelium* is a model organism for cellular motility and displays a wide range of responses to environmental clues. Positive and negative chemotaxis to a wide range of molecules have been documented but the cAMP response remains the most studied. Oxygen is an omnipresent molecule that can diffuse through membranes and thus constitute an odd signal to direct cell motility. Thus, the molecular mechanism behind aerotaxis in *Dd* remains a mystery but our present results further confirm that it is different from cAMP and classical G protein-coupled receptor based chemotaxis. They are summarized in Figure 6. In particular, aerotaxis functions only within a narrow range of ∼0.5% to 2% O_2_ at most. It has to be noted that, in water, atmospheric 2% O_2_ results into a concentration of ∼25 µM which might be too high to provide a spatial information for the molecular mechanism used to direct aerotactic motility. For comparison, cAMP acts in nanomolar range to orient cell motility (28). While starvation, like hypoxia, will ultimately results into lack of energy production in the form of ATP, Dd respond differentially to these challenges. When the cells are exposed to both simultaneously, they respond within minutes to very low O_2_ by aerotaxis to move out of the danger zone while chemotaxis will take hours to initiate as it requires the expression of a large group of specific genes. The directionality of aerotaxis, is far weaker than of cAMP chemotaxis (0.15 vs 0.9) as measured by the aerotactic or chemotactic indexes (29) and the cells lack the associated cell polarity (Figure 2F). It is therefore likely that the specialized actin polarization observed during chemotaxis is absent during aerotaxis. These results are compatible with a previous study that the G-beta protein is not involved in the sensing mechanism for aerotaxis contrary to cAMP chemotaxis (20).

**Figure 6:**
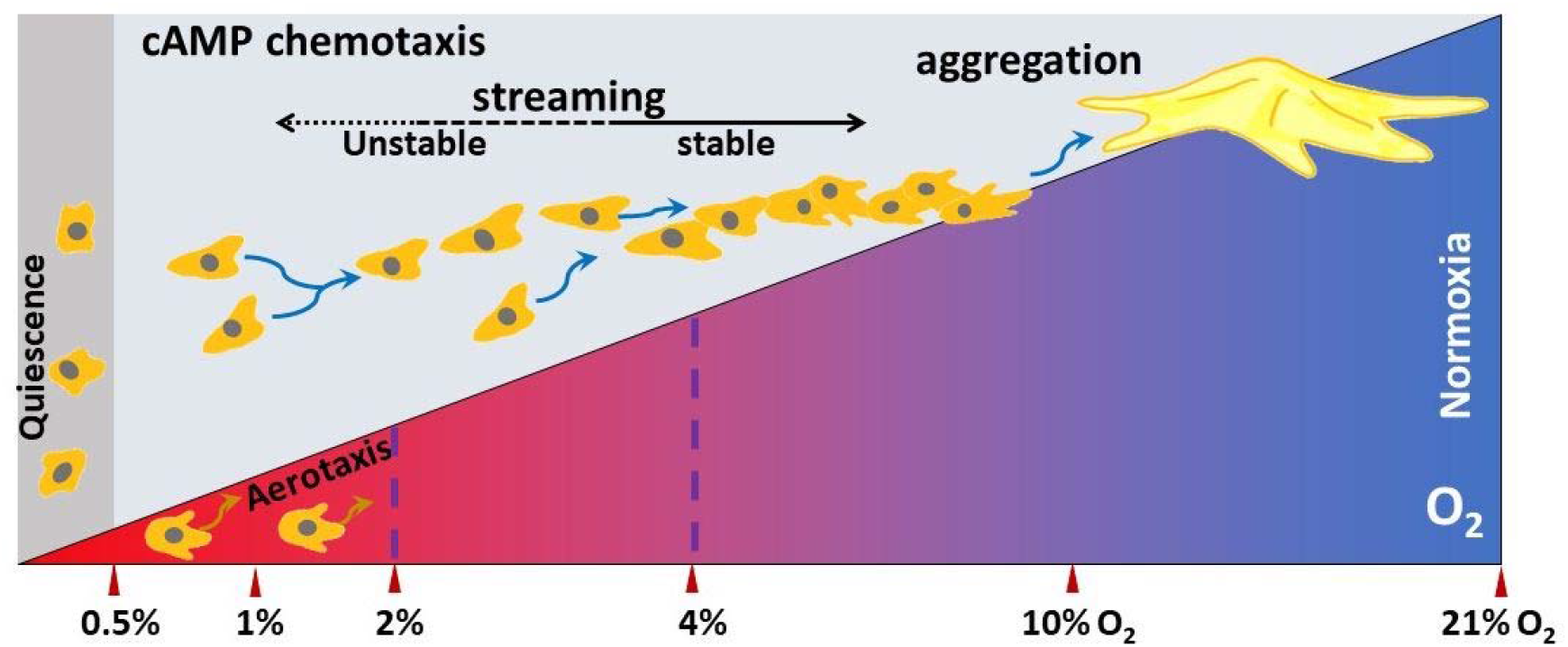
Schematic of the survival strategy of Dd. The aerotactic response enables cells to escape from hypoxic regions where aggregation cannot initiate. In severe hypoxia (up to 0.5% O_2_) near anoxia, Dd cells shrink, poorly adhere to the substratum and do not migrate. In a starvation buffer, between 0.5% and 2% O_2_, the cells exhibit aerotaxis, moving toward regions with higher O_2_ levels. They are also able to chemotax as optical density waves are visible but not to form aggregation streams. Streams start at ∼2% O_2_ but they quickly dissociate and hence aggregation centers are not stable. Stable aggregation centers form at more than 4% O_2_ and preferentially above 10% O_2_, thus attracting hypoxic cells to higher O_2_ level which allows them to aggregate and develop.

We have shown previously that low O_2_ does not prevent transcription nor translation as thousands of genes are specifically upregulated in response to this stress (29) and indeed, the transcription of aggregation competent genes is barely affected by hypoxia. However, our results show that hypoxia affects cAMP chemotaxis in a more subtle way: while cells can chemotax at 1% O_2_, they cannot establish a stable directionality but will follow the signal produced by cells that are forming an aggregation center at higher O_2_ level. Thus, the cells can use the efficiency of chemotaxis to move upward an O_2_ gradient in an O_2_ range where aerotaxis does not work. Our results also shows that it is possible to decoupled chemotactic response and formation of aggregation centers without gene mutation or using drugs, opening a new venue to study those two events and how they are normally coupled. We have shown that formation of an aggregation center requires the biosynthesis of a protein in an O_2_-rich environment. While stable aggregation centers do form at a uniform 4% O_2_, this might not correspond to an absolute threshold, but to a minimum, with higher O_2_ allowing better or faster formation of aggregation centers. This is consistent with the observation of only one aggregation center forming at 5% O_2_ in our gradient experiments while most occur at more than 15% O_2_ (Figure 3). It was proposed that initial aggregation centers correspond to random areas with an higher than average cell density that would produce more cAMP than nearby cells leading to further increase in cell density and preventing formation of other aggregation centers nearby (17). When O_2_ diffusion is limited, the increase in cell density will result in rapid depletion due to respiration. Dd appears to have evolved a fail-safe mechanism to prevent aggregation in a place where high cell density would be detrimental. Respiration in a non-vascular multicellular structure more than three cell deep usually results into anoxia leading to a necrotic core (reviewed in (30)). Such problem will be exacerbated if the outside O_2_ level is already low. The cycloheximide experiment shows that the formation of aggregation center requires the synthesis of one or more proteins that is limited by low O_2_. While the proteins identities are unknown, the fact that chemotaxis function at low O_2_ exclude the core proteins required for cAMP relay, cell polarity or motility. Some signaling pathways are known to modulate the cAMP chemotaxis (31) but do not fully match our results. The conditioned medium factor (CMF) is accumulated by growing cells then secreted upon starvation, acting as a quorum sensing factor to allow streaming only if the cell density is high enough (32) by limiting the activation of the cAMP signaling pathways downstream of CAR1 (33). Exposure to 1% O_2_ could result into a lack of secretion of CMF or act downstream but the resulting blockage is far stronger than what we observe: it would also prevent the accumulation of csaA mRNA (34) while we observed near normal level of induction. Furthermore, the high expression of the late streaming gene tgrC1 and the presence of bands in dark field at 1% O_2_ indicate that the cAMP relay is functioning sufficiently to induce gene expression and chemotactic motility. However, we cannot completely exclude that the CMF pathway or some of its components are involved in the prevention of streaming of starving cells in 1% O_2_.

It is not possible to determine if the mechanism behind the stream disaggregation observed with 2% O_2_ is a variant of the inhibition of streaming at 1% O_2_ or if it uses a completely different pathway. The second hypothesis seems more likely since with 2% O_2_, there are clear aggregation centers and streams are initially well formed with cell touching and following each other, meaning that cell-cell adhesion has been initiated, involving first CadA then CsaA proteins (35). The rapid disaggregation propagating along the stream involves breaking this cell-cell adhesion and some coordinated cell motility away from each other for some time. Such disaggregation of stream observed at 2% O_2_ could results from an over stimulated countin factor (CF) signaling pathway that is involve in breaking streams when the local cell density is too high (36). However, the classical over stimulation of this pathway observed in the smlA null mutant results in the formation of many tiny aggregates rather than the complete dissolution of the streams and aggregation centers (37,38). Furthermore, CF is a high molecular weight multiprotein complex secreted constantly by starving cells (39) that break streaming when reaching high local concentration within a stream. This appears incompatible with the fast-spreading wave of disaggregation we observed with 2% O_2_ as it affects simultaneously multiple streams that are not connected to each other and often more than 1 mm apart. It does seem possible that this disaggregation occurs through a modified CF pathway or reuse some of its components. Chemorepulsion like the one observed with AprA (40) or 8CPT-cAMP (41) might also be at play. AprA is normally secreted during cell growth and will induce peripheral cells to move away from a dense colony through the GrlH receptor (42), so it is unlikely AprA is directly involved, but another ligand might be used in hypoxia. 8CPT-cAMP is a synthetic analog of cAMP that act through CAR1 to trigger chemorepulsion (43). While 8CPT-cAMP is not known to occur naturally, it remains possible that *Dd* produces an analogous molecule during hypoxia since this chemorepulsion mechanism is clearly functional and efficient (44).

*Dictyostelium* has clearly evolved mechanisms to limit poor outcomes when facing multiple challenges simultaneously. As low environmental O_2_ levels were the norm till ∼600 million years ago, before the neoproterozoic oxygenation event and the generalization of multicellularity, it is likely that mechanisms to sense and avoid hypoxia appeared before the chemotactic system resulting into multicellular fruiting bodies (45). Indeed, the ancestral response to starvation in amoeba is the formation of unicellular microcyst that would limit the requirement to find a high O_2_ environment.

## Conclusion

Our work has uncovered unexpected interactions between two different common stress responses: to the lack of food or the lack of O_2_ necessary to metabolize food efficiently. In Dd, both stresses induce cell motility to reach a better outcome. While chemotaxis of Dd has been widely studied as a survival strategy, aerotaxis is a new phenomenon discovered in recent years. Our understanding is still limited but aerotaxis seems to enhance survival strategy by having a mechanism completely different from the known chemotaxis, contrary to electrotaxis (6). Aerotaxis is fast acting and doesn’t require *de novo* protein synthesis (18) but with a poor guidance efficiency (46), while chemotaxis is slower to initiate but more efficient (Figure 6). Surprisingly, hypoxia modulates the cAMP chemotactic response to avoid low O_2_ and instead promote aggregation at an O_2_ concentration higher than the functional range of aerotaxis. The undelying mechanisms remains unknown, but seem to influence both the position of aggregation centers and the stability of chemotactic streams to guide cells to an O_2_ rich environement, a useful strategy for a soil amoeba that form multicellular structure to increase its dispersal above ground. So the elucidation of those mechanism and how it affects chemotaxis could yield the discovery of a hidden mechanism in O_2_ response and survival strategy in eukaryotic cells, including human cell.

## Declarations

### Ethics approval and consent to participate

Not applicable.

### Consent for publication

All the authors have read and consented to the publication of this article.

### Availability of data and materials

The data presented in this study are available upon reasonable request to the corresponding authors.

### Authors’ contributions

Conceptualization: S.H., C.A. /Methodology: C.A., S.H., K.F. /Formal analysis: J-P.R, C.A., S.H., J.H. /Investigation: S.H., J.H., J-P.R, C.A. /Writing—original draft preparation: S.H., C.A./Writing—review and editing: S.H., C.A., J.H., J-P.R., K.F. /Visualization: J-P.R., S.H. /Funding acquisition: J-P.R., S.H., K.F.

## Acknowledgements

We thank Sandra Raymond for technical help cell in culture,RT-qPCR measurements and routine videomicroscopy. Some exploratory experiments were performed by first year Life science Master degree students: Inès Taïeb, Léa Bouchet and Tély Pandaure. We thank Pr Satochi Sawai for insightful discussion and comments. Part of the work was carried out under the Collaborative Research Project of the Institute of Fluid Science, Tohoku University.

## Supplementary Figures and Movies

**Supplementary Figure S1:**
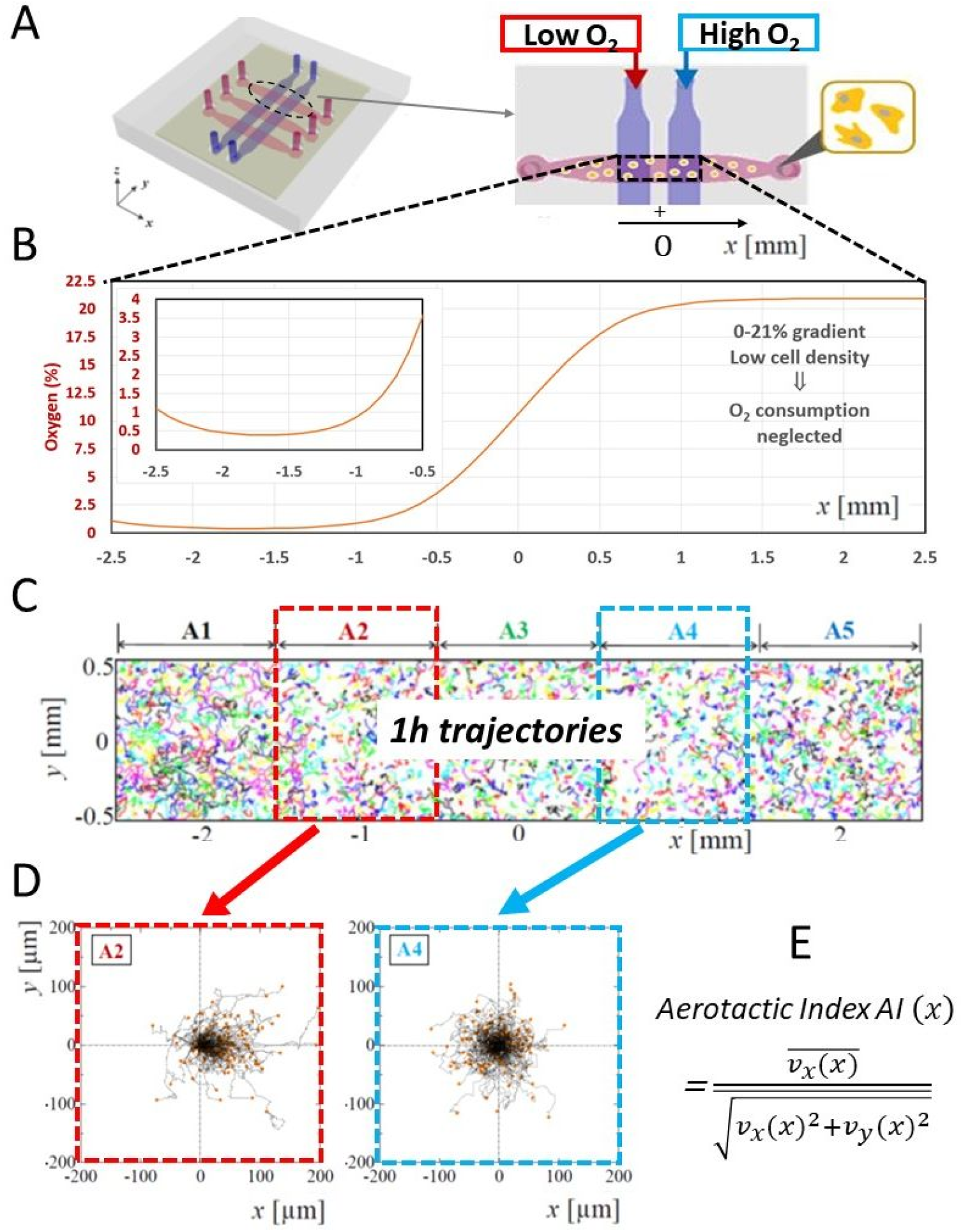
Microfluidic device to measure the cells’ aerotactic index. A) Schematic of the device: The two gas channels (blue) were located vertically above and perpendicular to the three media channels with cells (pink). In the experiments presented here, pure nitrogen (0% O_2_) and air (21% O_2_) was supplied into the left and right gas channels, respectively, to generate an O_2_ concentration gradient. B) The O_2_ profile is computed using COMSOL Multiphysics taking into account cell comsumption that reduces the O_2_ level when density is high. The zoom shows the region under the gas channel where Dd cells are aerotactic. C) Experimental trajectories of AX2 cells in HL5 medium lasting 1 h recorded between 1 h and 2 h after establishing the gradient. D) Trajectories of the cells in the areas A2 (0.4-3.4% O_2_ region) and A4 (17.5-20.5% O_2_ region), setting their initial positions at the origin. E) Definition of the aerotactic index as the ratio of the mean of the cell migration velocity along the *x* axis parallel to the media channel over the mean migration speed. The value was measured at 1-min intervals.

**Figure S2:**
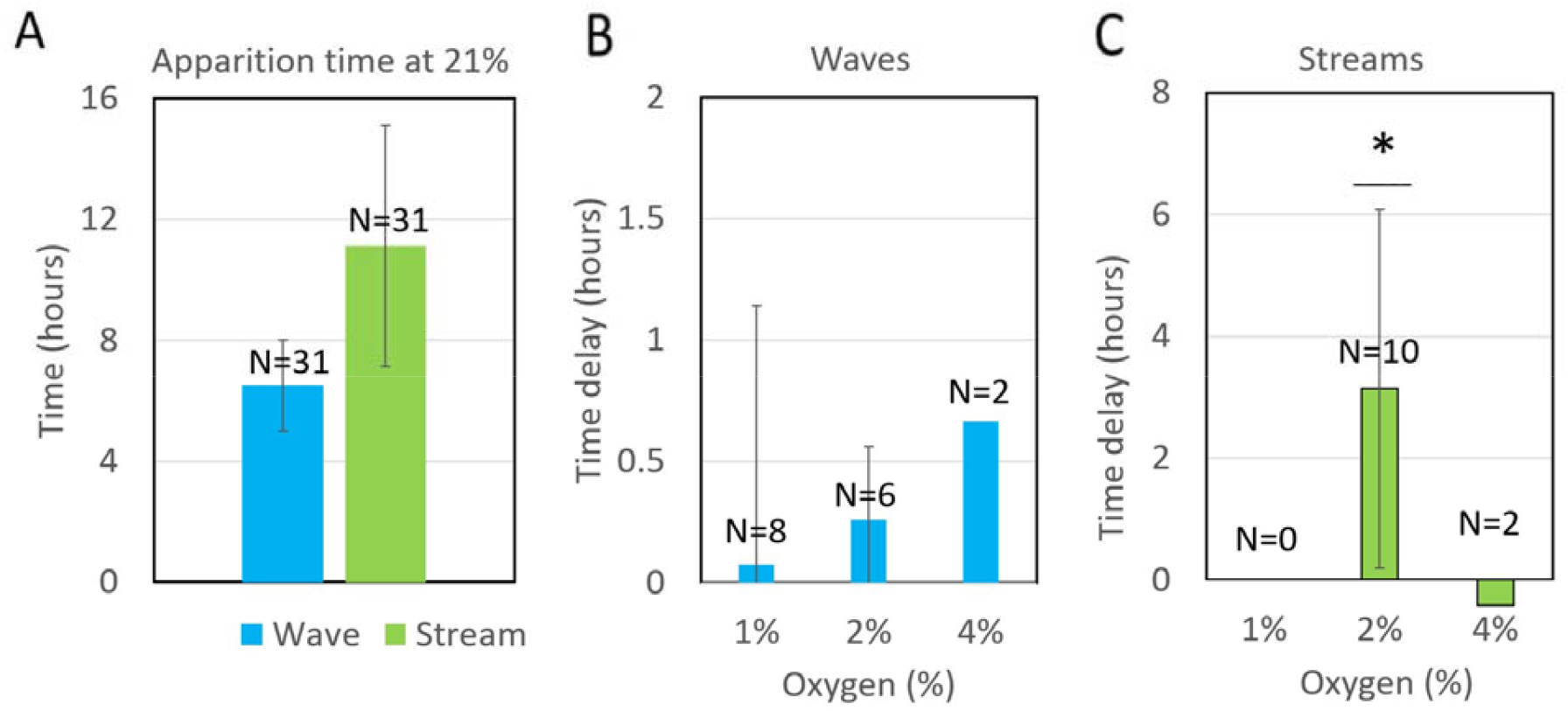
A) Mean apparition time of cAMP induced waves and streams in the control experiments at 21% O_2_. B-C) Delays in wave and streaming apparition between a given hypoxic condition and the normoxic control of the day. For each condition the number of independent experiment *N* is indicated. The streaming delay at 2% O_2_ is significantly different from 0 (*p* = 0.030, one-sample t-test between the condition and the expected 0 difference value).

**Supplementary Movie M1**. Development of starved Dd in a 0-21% O_2_ gradient generated by a microfluidic device. Cells start to aggregate on the device top or bottom borders mostly on the left side where O_2_ is higher than 10% O_2_. Time is indicated as h:min. The bottom part shows the O_2_ profile calculated using COMSOL Multiphysics.

**Supplementary Movie M2**. Starvation response under homogenous 1% O_2_. (right column) as compared to normoxia (left column). Raw dark field images are presented on the top raw and the substraction image using a 10-min interval between substracted images is presented on the lower raw. The scale bar indicated on the bottom right corresponds to 1 mm while the time from starvation (hour:minutes) is indicated on the top right corner. Within 12h40, the control condition shows aggregation streams while the 1% O_2_ condition (middle) present only disorganized waves during 24 h. After 24 h, the sample was reoxygenated and within 3 h, the previously hypoxic sample starts to stream and aggregate (right).

**Supplementary Movie M3**. Starvation response under homogenous 2% O_2_. (right column) as compared to normoxia (left column). Raw dark field images are presented on the top raw and the substraction image using a 10-min interval between substracted images is presented on the lower raw. The scale bar indicated on the bottom right corresponds to 1 mm while the time from starvation (hour: minutes) is indicated on the top right corner. At 2% O_2_, the sample starts to stream with some delay with the control but streams quickly dissociate after their formation and aggregates do not form.

**Supplementary Movie M4**. Starvation response under homogenous 4% O_2_. (right column) as compared to normoxia (left column). Raw dark field images are presented on the top raw and the substraction image using a 10-min interval between substracted images is presented on the lower raw. The scale bar indicated on the bottom right corresponds to 1mm while the time from starvation (hour: minutes) is indicated on the top right corner. At 4% O_2_ streaming do occur at similar time than control and are mostly stable but continue for longer time before forming stable aggregates.

**Supplementary Movie M5**. Reoxygenation of starved cells that were initially incubated 18h at 1% O_2_ in presence (right) or absence (DMSO, left) of cycloheximide. The control at 21% O_2_ is also presented on left panel. Time of the picture are indicated on the top right, the scales bar correspond to 2 mm.

